## Supplementary Table 1 for "Functional genomics reveals strain-specific genetic requirements conferring hypoxic growth in *Mycobacterium intracellulare*"

Supplemental Table 1: Genotypes of study strains and the strains used for each experiment.

| Strain | Genotype | TnSeq in *vitro* | TnSeq in mice | Hypoxic growth assay | Knockdown experiment |
| --- | --- | --- | --- | --- | --- |
| ATCC13950 | TMI | Done |  | Done | Done |
| M.i.198 | TMI | Done | Done | Done | Done |
| M.i.27 | TMI | Done | Done | Done | Done |
| M018 | TMI | Done |  | Done | Done |
| M003 | MP-MIP | Done |  | Done | Done |
| MOTT64 | MP-MIP | Done |  | Done |  |
| M019 | MP-MIP | Done |  | Done | Done |
| M001 | MP-MIP | Done |  | Done |  |
| M021 | MP-MIP | Done |  | Done | Done |
| M005 | TMI |  |  | Done |  |
| M016 | TMI |  |  | Done |  |

TMI: Typical *M. intracellulare*, MP-MIP: *M. paraintracellulare-M. indicus pranii*
