## Supplementary Information for "Functional genomics reveals strain-specific genetic requirements conferring hypoxic growth in *Mycobacterium intracellulare*"

#### Supplementary Figure 1

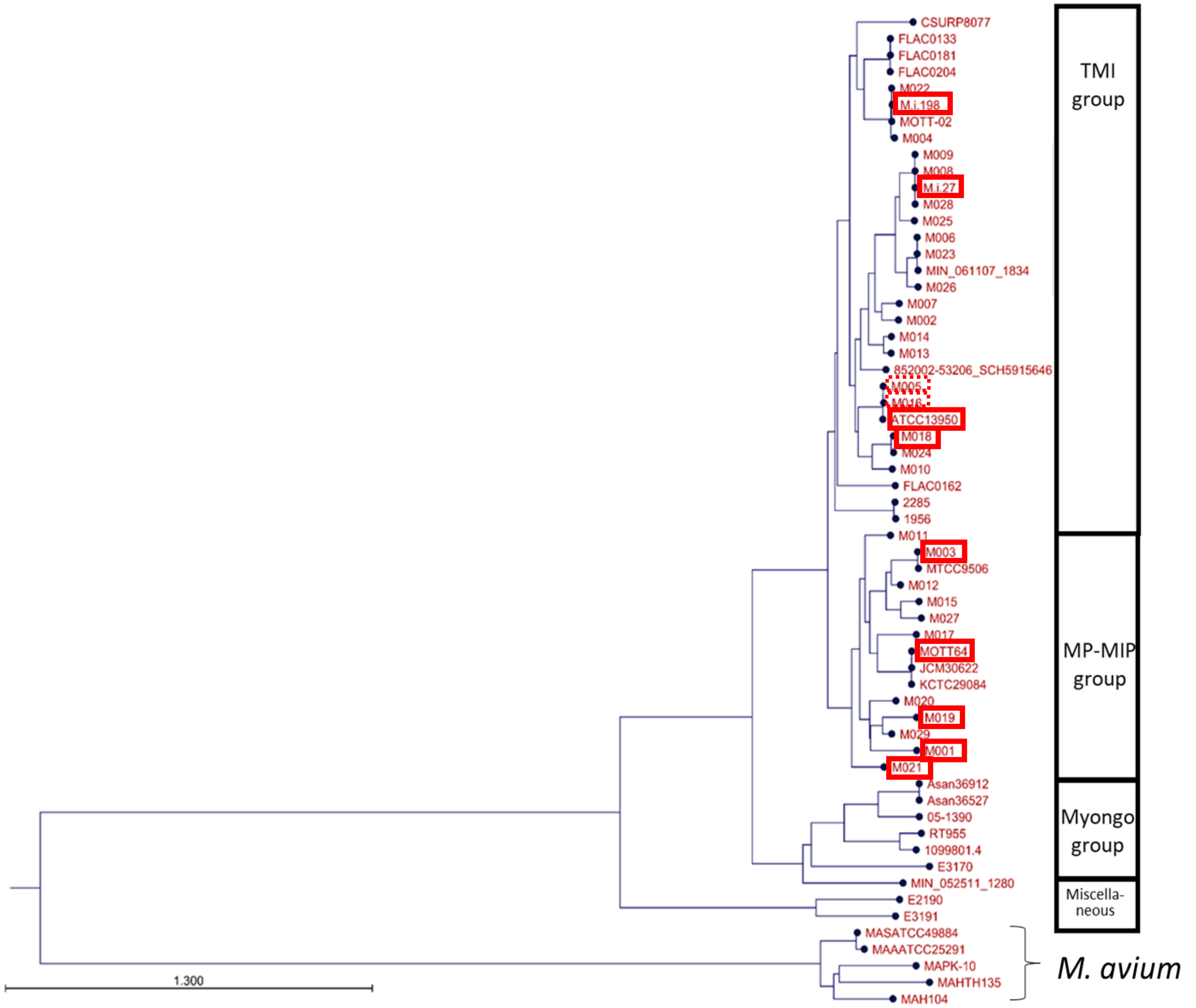


**Supplementary Figure 1.** Subject *M. intracellulare* strains in this study. The phylogenetic tree represents the global genetic diversity of the *M. intracellulare* strains by our previous study[5]. The subject strains were squared in red. As for strains squared in bold line, four strains including the type strain ATCC13950 were selected from TMI (typical *M. intracellulare*) group, and 5 strains were selected from MP-MIP (*M. paraintracellulare-M. indicus pranii*) group in this TnSeq study. Two strains squared in dashed line are closely related neighbors of ATCC13950, which were additionally enrolled in the hypoxic growth assay (as explained in **Supplementary Table 1**).

**Supplementary Figure 2**

| | |
|---|---|
| Enrichment Score (ES) | 0.37941608 |
| Normalized Enrichment Score (NES) | 1.8299032 |
| Nominal p-value | 0.055555556 |
| FDR q-value | 0.055555556 |
| FWER p-Value | 0.001 |

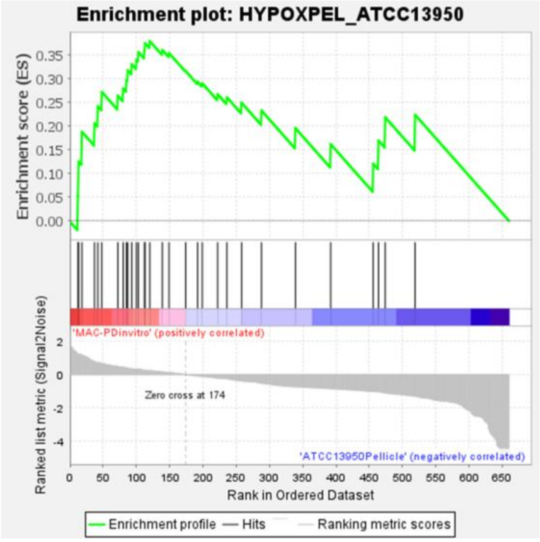


**Supplementary Figure 2.** Summary of the gene set enrichment analysis (GSEA) results. The dataset of strain-dependent/accessory essential or growth-defect-associated genes was statistically tested to be enriched with the gene sets required for hypoxic pellicle formation observed in ATCC13950. **a** Statistics data for the gene set enrichment analysis. **b** Data of enrichment plots. On X-axis, the left direction indicates positive correlation with the gene sets required for hypoxic pellicle formation observed in ATCC13950, and the right direction indicates negative correlation with the gene sets required for hypoxic pellicle formation observed in ATCC13950. Only 31 genes were ranked because the rest of 144 genes were not present in the gene sets of strain-dependent or accessory essential genes.

The ranking of the genes in the gene sets required for hypoxic pellicle formation observed in ATCC13950 is shown in **Supplementary Table 6**.

#### Supplementary Figure 3

**a** *M. intracellualre* strains for infection

- Hypervirulence strain M.i.198Tn mutant library strain : $3.25 \times 10^8$ CFUs
- Intermediate virulence strain M.i.27Tn mutant library strain : $5.93 \times 10^7$ CFUs

Six-week old female C57BL/6JJcl mice were infected with these strains.

The experimental method of mouse infection was based on our previous publication[22].

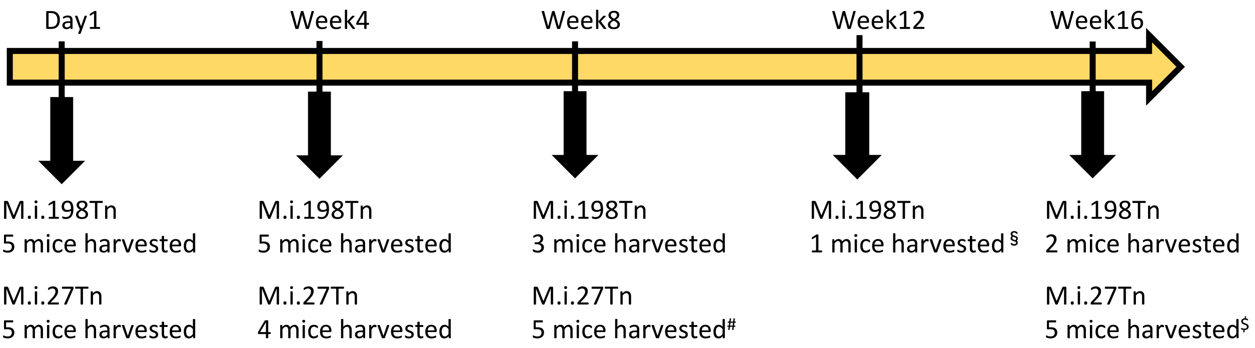


**b**

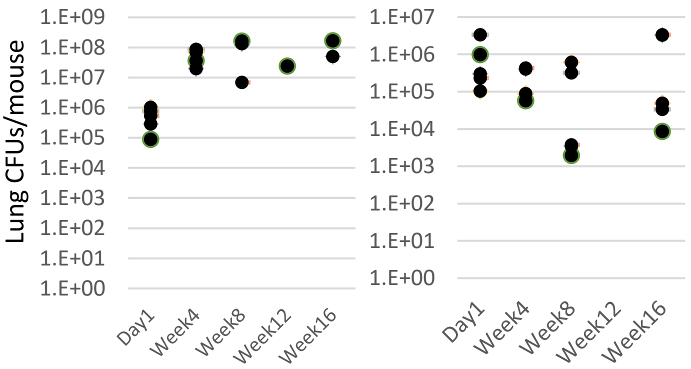


**c**

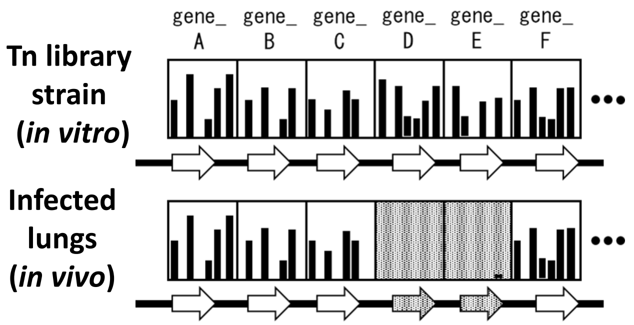


**d**

| | Day1 | Week4 | Week8 | Week16 |
|---|---|---|---|---|
| M.i.198Tn | 324 | 429 | 215 | 257 |
| M.i.27Tn | 213 | 246 | 210 | 442 |

**Supplementary Figure 3.** Mouse infection experiment for TnSeq **a** Schematic representation of mouse infection experiment. #Three mice were excluded from TnSeq because of the number of harvested colonies was < 1 × $10^4$/mouse. §Three samples of M.i.198Tn at week 12 were grouped into M.i.198Tn at week 16 to represent the chronic phase of infection. $Three mice were excluded from TnSeq because the number of harvested colonies was only $10^3$–$10^4$/mouse. **b** Time-course changes in colony-forming units (CFUs) in mouse lungs after infection with M.i.198Tn or M.i.27Tn. Data on CFUs are from one biological experiment ($N$ = 1). Each dot represents an individual mouse. **c** Resampling assay to detect significantly reduced Tn insertion reads in infected mouse lungs compared with before infection. The data of time-course changes in colony-forming units (CFUs) in mouse lungs after infection with wild type strains of M.i.198 or M.i.27 were published previously[22]. **d** Number of genes showing significantly reduced Tn insertion reads in the infected mouse lungs compared with before infection according to a resampling analysis. As described above, the samples of M.i.198Tn in week 16 were derived from two mice killed in week 16 and one mouse killed in week 12.

#### Supplementary Figure 4

| | M.i.27 | M.i.198 |
|---|---|---|
| Enrichment Score (ES) | 0.6164138 | 0.39179814 |
| Normalized Enrichment Score (NES) | 1.8604097 | 1.3387576 |
| Nominal p-value | 0.0 | 0.01629328 |
| FDR q-value | 0.0 | 0.01629328 |
| FWER p-Value | 0.0 | 0.016 |

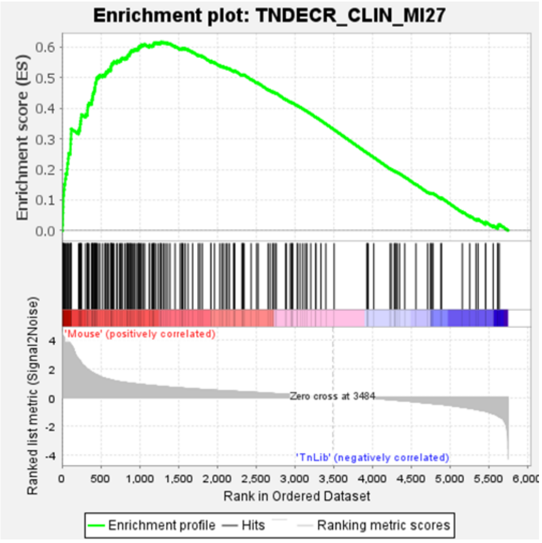


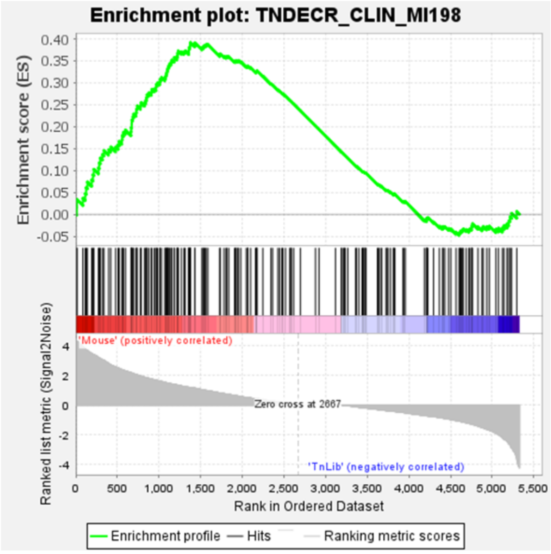


**Supplementary Figure 4.** Summary of the gene set enrichment analysis (GSEA) results. The dataset of fitness costs for infection in mouse lungs in Tn mutant libraries of M.i.27 and M.i.198 from day 1 to week 16 of infection was statistically tested to be enriched with the gene sets required for hypoxic pellicle formation observed in ATCC13950 and with those showing increased genetic requirements observed in the MAC-PD clinical strains. **a** Statistics data for the gene set enrichment analysis. **b** Data of enrichment plots. On X-axis, the left direction indicates positive correlation with the conditions after infection in mouse lungs, and the right direction indicates positive correlation with the conditions outside of mouse lungs (i.e. negative correlation with the conditions after infection in mouse lungs). The ranking of the genes in the gene sets required for hypoxic pellicle formation observed in ATCC13950 and those showing increased genetic requirements observed in the MAC-PD clinical strains is shown in **Supplementary Table 12**.

#### Supplementary Figure 5

| | M.i.27_Day 1 to Week 16 | M.i.27_Week 4 to Week16 | M.i.198_Day 1 to Week16 | M.i.198_Week 4 to Week16 |
|---|---|---|---|---|
| Enrichment Score (ES) | 0.6164929 | 0.61192846 | 0.38254246 | 0.45932388 |
| Normalized Enrichment Score (NES) | 1.8427752 | 1.82273 | 1.3169765 | 1.4549882 |
| Nominal p-value | 0.0 | 0.0 | 0.026557712 | 0.0020120724 |
| FDR q-value | 0.0 | 0.0 | 0.026557712 | 0.0020120724 |
| FWER p-Value | 0.0 | 0.0 | 0.026 | 0.002 |

M.i.27_Day 1 to Week 16

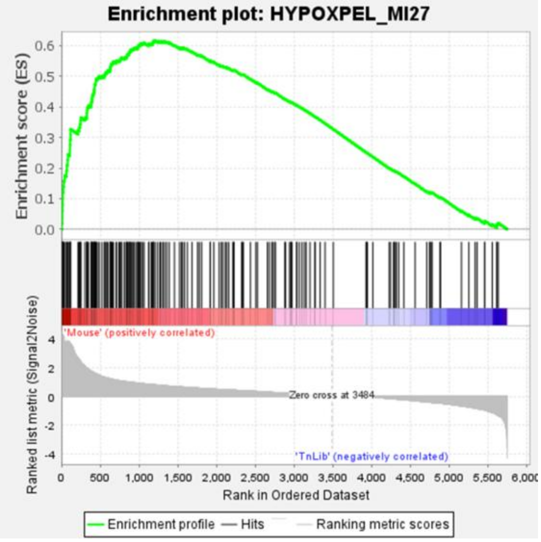


M.i.27_Week 4 to Week 16

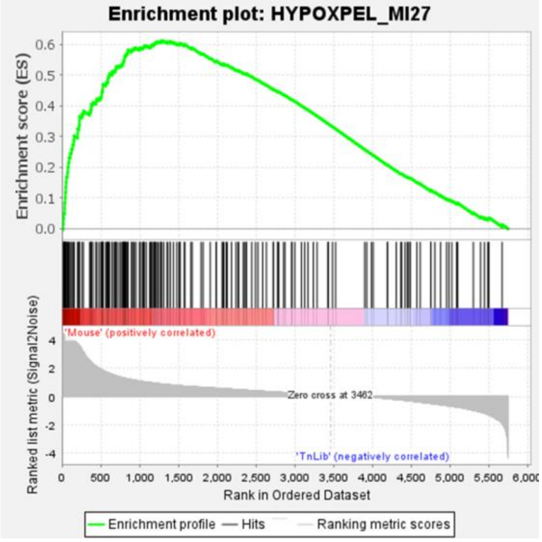


M.i.198_Day 1 to Week 16

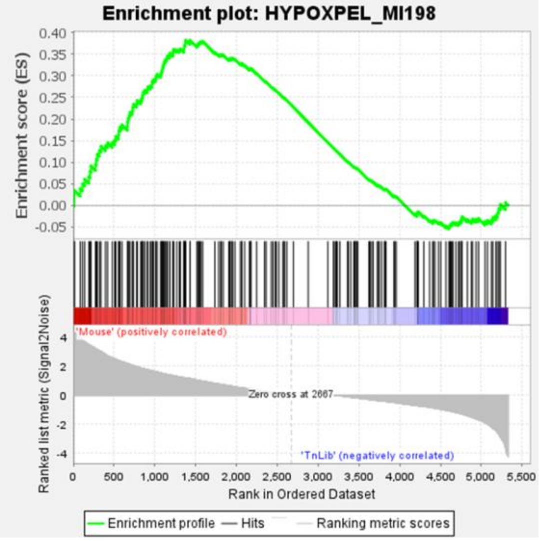


M.i.198_Week 4 to Week 16

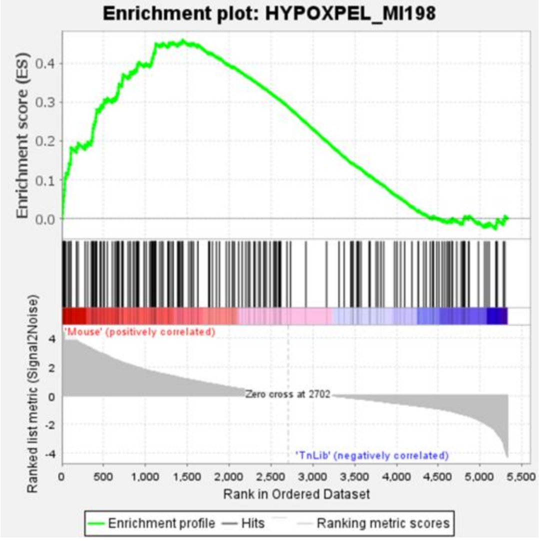

**Supplementary Figure 5.** Summary of the gene set enrichment analysis (GSEA) results. The dataset of fitness costs for infection in mouse lungs in Tn mutant libraries of M.i.27 and M.i.198 was statistically tested to be enriched with the gene sets required for hypoxic pellicle formation observed in ATCC13950. **a** Statistics data for the gene set enrichment analysis. **b** Data of enrichment plots. On X-axis, the left direction indicates positive correlation with the conditions after infection in mouse lungs, and the right direction indicates positive correlation with the conditions outside of mouse lungs (i.e. negative correlation with the conditions after infection in mouse lungs). The ranking of the genes in the gene sets required for hypoxic pellicle formation observed in ATCC13950 is shown in **Supplementary Table 16**.

### Supplementary Figure 6

| | M.i.27_Day 1 to Week 16 | M.i.27_Week 4 to Week16 | M.i.198_Day 1 to Week16 | M.i.198_Week 4 to Week16 |
|---|---|---|---|---|
| Enrichment Score (ES) | 0.6494766 | 0.64232165 | 0.50276554 | 0.65003544 |
| Normalized Enrichment Score (NES) | 1.9255978 | 1.8758564 | 2.3044288 | 2.034254 |
| Nominal p-value | 0.0 | 0.0 | 0.0 | 0.0 |
| FDR q-value | 0.0 | 0.0 | 0.0 | 0.0 |
| FWER p-Value | 0.0 | 0.0 | 0.0 | 0.0 |

M.i.27_Day 1 to Week 16

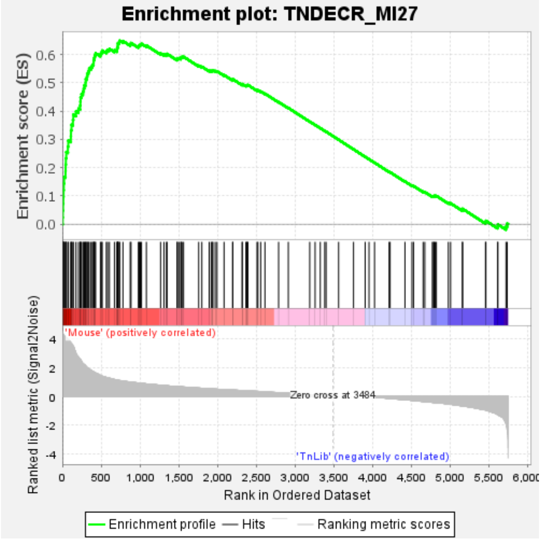


M.i.27_Week 4 to Week 16

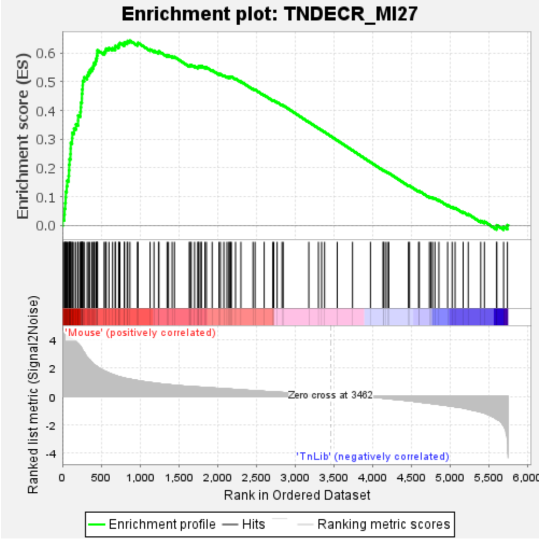


M.i.198_Day 1 to Week 16

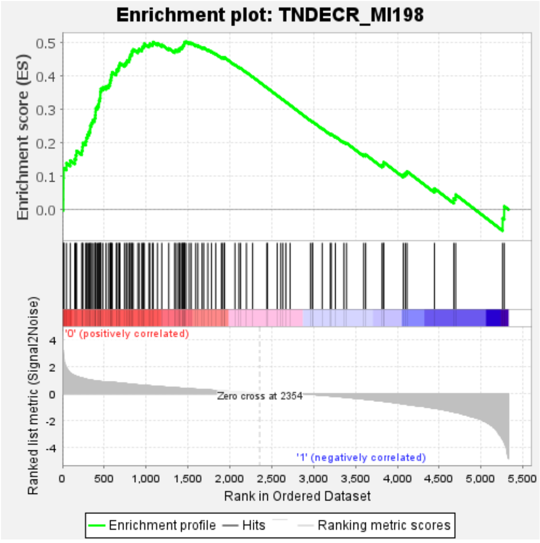


M.i.198_Week 4 to Week 16

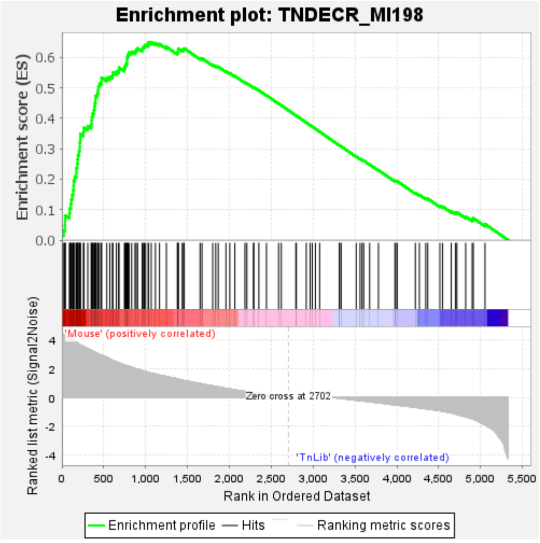

**Supplementary Figure 6.** Summary of the gene set enrichment analysis (GSEA) results. The dataset of fitness costs for infection in mouse lungs in Tn mutant libraries of M.i.27 and M.i.198 was statistically tested to be enriched with those showing increased genetic requirements observed in the MAC-PD clinical strains. **a** Statistics data for the gene set enrichment analysis. **b** Data of enrichment plots. On X-axis, the left direction indicates positive correlation with the conditions after infection in mouse lungs, and the right direction indicates positive correlation with the conditions outside of mouse lungs (i.e. negative correlation with the conditions after infection in mouse lungs). The ranking of the genes showing increased genetic requirements observed in the MAC-PD clinical strains is shown in **Supplementary Table 17**.

#### Supplementary Figure 7

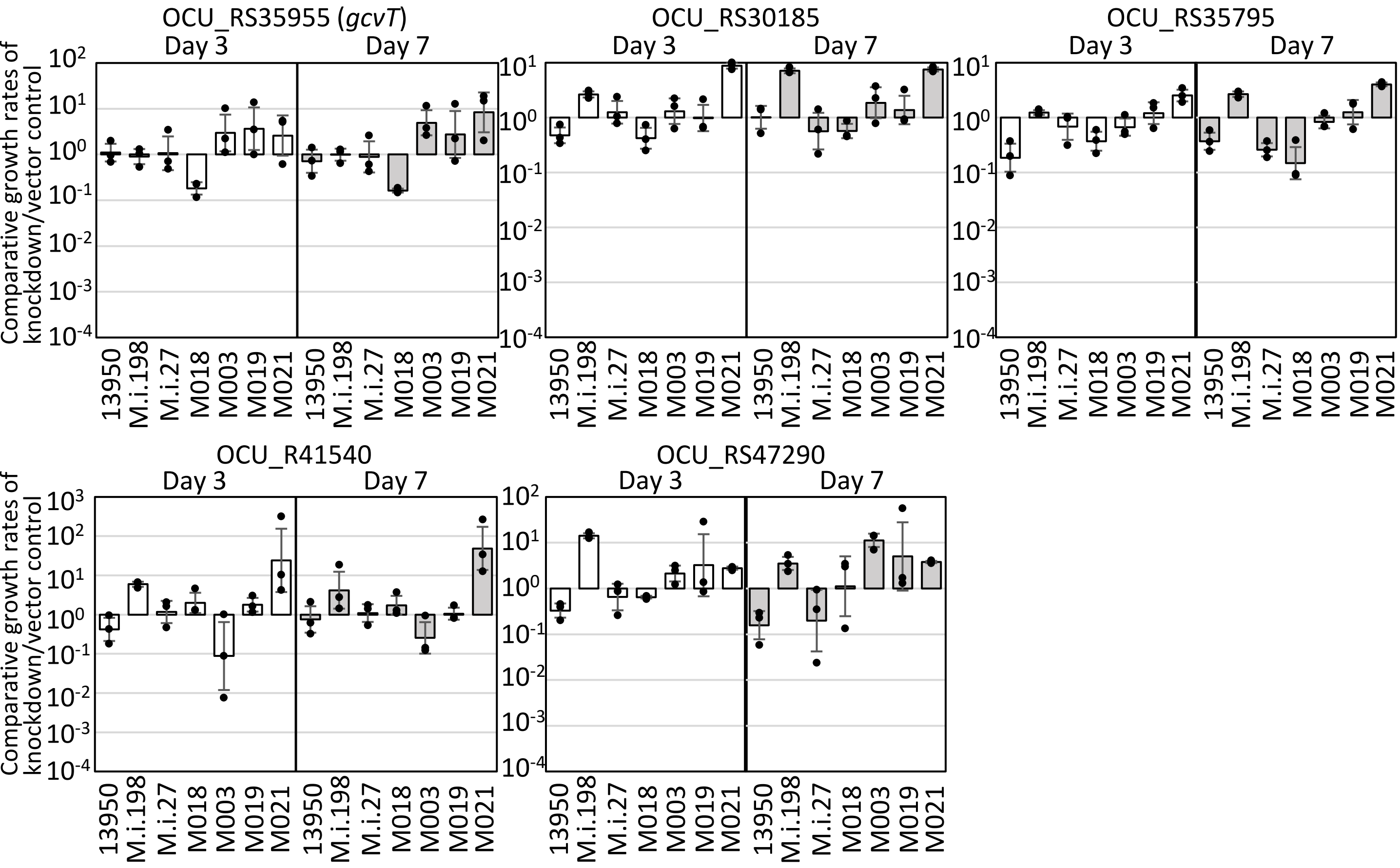


**Supplementary Figure 7.** Comparative growth rates of the knockdown strains relative to those of the vector control strains in several accessory and strain-dependent essential or growth-defect-associated genes. Data are shown as the means ± SD of triplicate experiments. Data from one experiment representative of two independent experiments (*N* = 2) are shown.

#### Supplementary Figure 8

**a**

| | | |
|---|---|---|
| KDCn#01 | OCU_RS38100 | Malate synthase G GlcB |
| KDCn#03 | OCU_RS40185 | Enoyl[acyl-carrier –prtotein] reductase FabI (InhA) |
| KDCn#05 | OCU_RS25020 | DNA topoisomerase subunit B GyrB |
| KDCn#07 | OCU_RS26150 | Arabinosyltransferase EmbB |

**b**

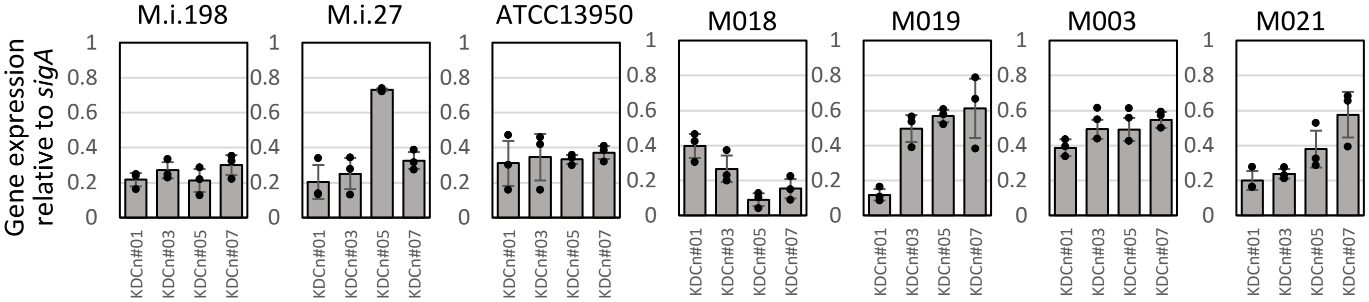


**c**

| | | |
|---|---|---|
| KD-1 | OCU_RS30540 | Fructose bisphosphatase GlpX |
| KD-2 | OCU_RS38345 | Type VII secretion protein EccC |
| KD-3 | OCU_RS48660 | Phosphoenolpyruvate carboxykinase PckA |
| KD-4 | OCU_RS30185 | Ppx/GppA family phosphatase |
| KD-5 | OCU_RS35795 | Bifunctional RNase H/acid phosphatase |
| KD-6 | OCU_RS35955 | Glycine cleavage system GcvT |
| KD-7 | OCU_RS38275 | type VII secretion-associated serine protease mycosin MycP5 |
| KD-8 | OCU_RS40305 | Cysteine desulfurase Csd |
| KD-9 | OCU_RS41540 | Class I SAM-dependent RNA methyltransferase |
| KD-10 | OCU_RS47290 | Hypothetical protein |

**d**

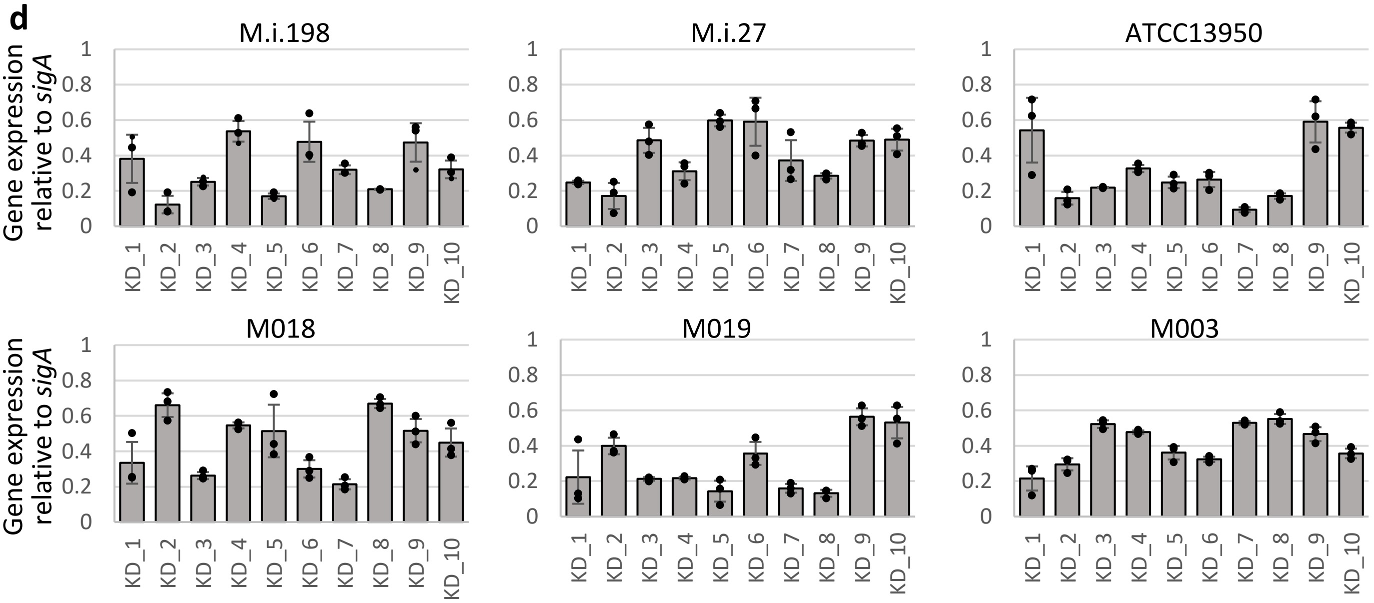


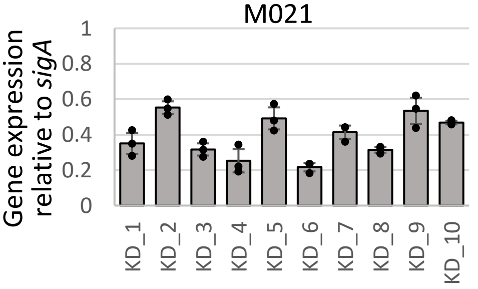

**Supplementary Figure 8.** qRT-PCR validation of the suppression of gene expression in knockdown strains of universal and strain-dependent/ accessory essential or growth-defect-associated genes. **a** List of target universal essential or growth-defect-associated genes analyzed with qRT-PCR. **b** Levels of gene expression of the target universal essential or growth-defect-associated genes in knockdown strains compared with those in control strains not expressing guide RNA. **c** List of target strain-dependent/ accessory essential or growth-defect-associated genes analyzed with qRT-PCR. **d** Levels of gene expression of the target strain-dependent/ accessory essential or growth-defect-associated genes in knockdown strains compared with those in control strains not expressing guide RNA. Aliquots (10 mL) of log-phase bacterial cultures were prepared and the expression of guide RNA was induced by the addition of aTc on day 0 and day 2 (final concentrations of 50 ng/ml for ATCC13950 and M.i.27; 200 ng/ml for M.i.198, M018, M019, M003, and M021). The bacteria were harvested, RNA extracted, and qRT-PCR performed to evaluate the gene expression levels. Gene expression data are the means ± SD of triplicate experiments. Data from one experiment representative of two independent experiments (*N* = 2) are shown.

#### Supplementary Figure 9

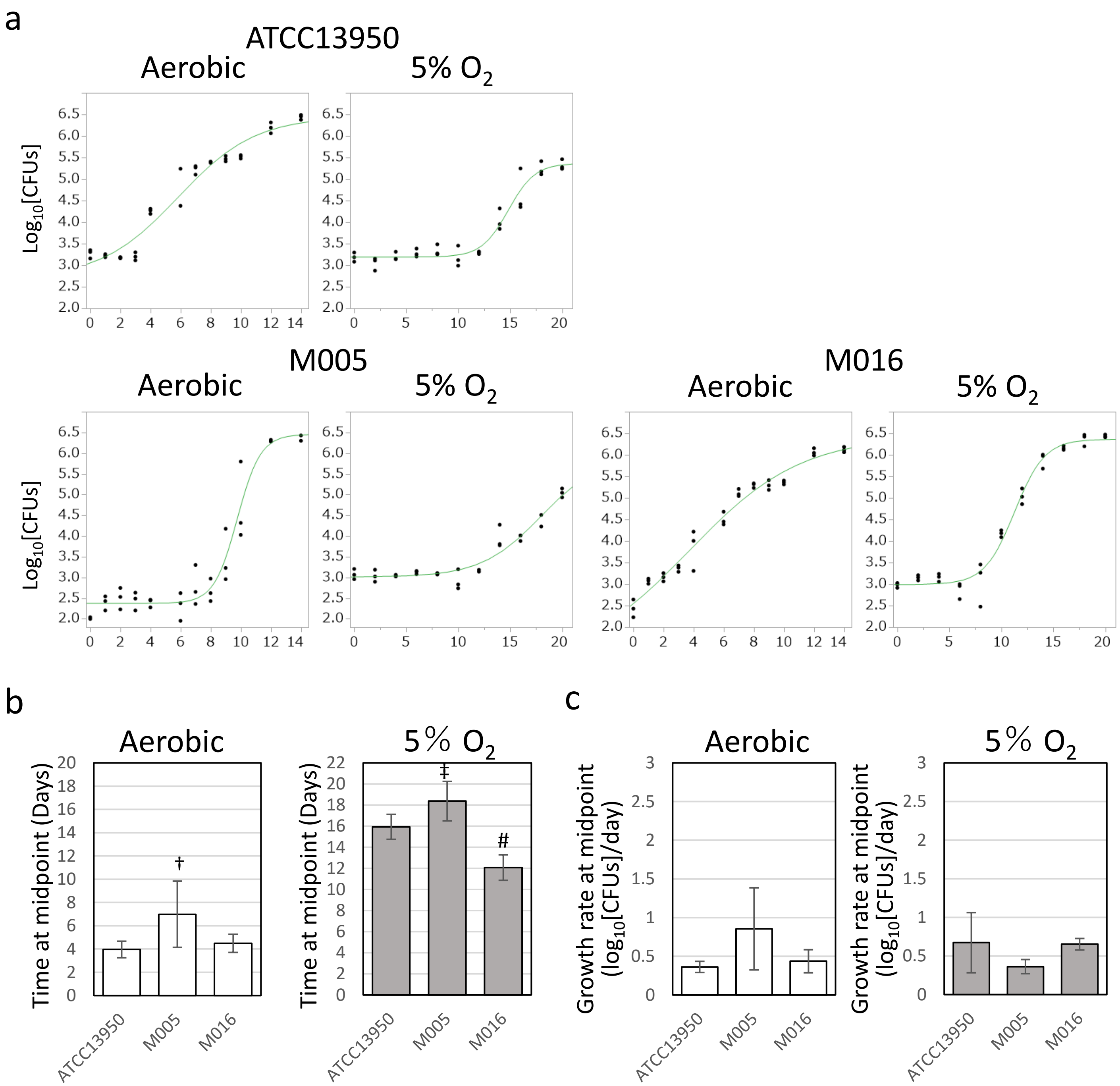

**Supplementary Figure 9. Data of growth curve in ATCC13950 and its neighbor clinical *M. intracellulare* strains M005 and M016. a** Representative data on the growth curves of ATCC13950, M005 and M016 under aerobic and 5% oxygen conditions. The assay was performed three times in 96-well plates containing 250 µl of broth medium per well, in triplicate. Data are represented as CFUs in 4 µl sample at each timepoint. Data from one experiment representative of three independent experiments (*N* = 3) are shown. **b** Comparison of the time at the inflection point (midpoint) on the sigmoid growth curve between the clinical *M. intracellulare* strains M005 and M016 and the type strain ATCC13950. $^{\#}$significantly earlier than a hypoxic culture of ATCC13950; $^{\dagger}$significantly later than an aerobic culture of ATCC13950; $^{\ddagger}$significantly later than a hypoxic culture of ATCC13950. **c** Comparison of the logarithmic growth rate at midpoint between the clinical *M. intracellulare* strains M005 and M016 and the type strain ATCC13950. Open bars: aerobic; closed bars: 5% $O_2$. Data are shown as the means ± SD of triplicate experiments.

**Supplementary Table 1.** Genotypes of study strains and the strains used for each experiment.

**Supplementary Table 2.** Number of reads obtained and TA coverage in each replicate of Tn mutant library strains. Data for ATCC13950 are from our previous study[10].

**Supplementary Table 3.** Number of essential, growth-defect-associated, nonessential, and growth-advantage-associated genes detected with an HMM analysis in each *M. intracellulare* strain. Data for ATCC13950 are from our previous study[10].

**Supplementary Table 4.** List of genes identified as universal essential or growth-defect-associated among the nine *M. intracellulare* strains analyzed in this study. ES: essential, GD: growth-defect-associated.

**Supplementary Table 5.** List of genes identified as accessory essential or growth-defect-associated in the accessory genome and strain-dependent essential or growth-defect-associated in the core genes of *M. intracellulare*. "Hit" represents the genes identified as essential or growth-defect-associated in each strain. The data of pan-genome were referred to our previous study[5].

**Supplementary Table 6.** Result of the gene set enrichment analysis (GSEA) in strain-dependent/accessory essential or growth-defect-associated genes. Genes required for hypoxic pellicle formation observed in ATCC13950 were ordered by their position in the ranked list of genes. Only 31 genes were ranked because the rest of 144 genes were not present in the gene set of strain-dependent or accessory essential genes.

**Supplementary Table 7.** Classification of the genes showing increased genetic requirements in the clinical MAC-PD strains compared to ATCC13950, and the genes of gluconeogenesis, fructose-1,6-bisphosphatase *glpX,* with respect to the core and accessory genomes.

**Supplementary Table 8**. List of genes showing significantly increased or reduced Tn insertion reads in the clinical *M. intracellulare* strains compared with ATCC13950 by a resampling analysis, with information of the genetic requirements detected by an HMM analysis.

**Supplementary Table 9.** Number of Tn insertion reads in samples from infected mouse lungs. Individual mice are represented as A–E after the sampling time points.

**Supplementary Table 10.** List of genes identified with a resampling analysis in mouse lungs infected with M.i.198 Tn mutant library strains.

**Supplementary Table 11.** List of genes identified with a resampling analysis in mouse lungs infected with M.i.27 Tn mutant library strains.

**Supplementary Table 12.** List of genes required for mouse lung infection that are also required for hypoxic pellicle formation by ATCC13950.

**Supplementary Table 13.** List of genes required for mouse lung infection at all time points from Day 1 to Week 16, and  from Week 4 to Week 16.

**Supplementary Table 14.** List of genes required for mouse lung infection at all time points from Day 1 to Week 16, and from Week 4 to Week 16, as well as required for hypoxic pellicle formation in ATCC13950.

**Supplementary Table 15.** Result of the gene set enrichment analysis (GSEA). Genes in the gene sets required for hypoxic pellicle formation observed in ATCC13950 and those showing increased genetic requirements observed in the clinical MAC-PD strains were ordered by their position in the ranked list of genes.

**Supplementary Table 16.** Result of the gene set enrichment analysis (GSEA) in mouse TnSeq datasets. Genes required for hypoxic pellicle formation observed in ATCC13950 were ordered by their position in the ranked list of genes.

**Supplementary Table 17.** Result of the gene set enrichment analysis (GSEA) in mouse TnSeq datasets. Genes showing increased genetic requirements observed in the clinical MAC-PD strains were ordered by their position in the ranked list of genes.

**Supplementary Table 18.** Correlation analysis between the fitness costs ($log_2FC$), efficiency of knockdown (qRT-PCR) and growth suppression in the knockdown strains of strain-dependent/accessory essential or growth-defect-associated genes.

**Supplementary Table 19.** The raw data of CFUs used for drawing growth curves in Fig. 8a.

**Supplementary Table 20.** Oligonucleotides and primers used in this study.
